# p53 dynamically directs TFIID assembly on target gene promoters

**DOI:** 10.1101/083014

**Authors:** R. A. Coleman, Z. Qiao, S. K. Singh, C. S. Peng, M. Cianfrocco, Z. Zhang, A. Piasecka, H. Aldeborgh, G. Basishvili, W. L. Liu

## Abstract

The p53 tumor suppressor protein is a central regulator that turns on vast gene networks to maintain cellular integrity upon various stimuli. p53 activates transcription initiation in part by aiding recruitment of TFIID to the promoter. However, the precise means by which p53 dynamically interacts with TFIID to facilitate assembly on target gene promoters remains elusive. To address this key question, we have undertaken an integrated approach involving single molecule fluorescence microscopy, single particle cryo-electron microscopy, and biochemistry. Our real-time single molecule imaging demonstrates that TFIID alone binds poorly to native p53 target promoters. p53 unlocks TFIID’s ability to bind DNA by increasing TFIID contacts with both the core promoter and a region surrounding p53’s response element (RE). Analysis of single molecule dissociation kinetics reveals that TFIID interacts with promoters via transient and prolonged DNA binding modes that are each regulated by p53. Importantly, our structural work reveals that TFIID’s conversion from a canonical form to a rearranged DNA-binding conformation is enhanced in the presence of DNA and p53. Notably, TFIID’s interaction with DNA induces p53 to rapidly dissociate, effectively liberating the RE on the promoter. Collectively, these findings indicate that p53 dynamically escorts and loads the basal transcription machinery onto its target promoters.

## Introduction

More than 50% of cancer patients harbor p53 mutations, highlighting the essential role of this protein in tumor suppression (1). To properly maintain genomic stability, p53 induces vast gene networks involved in diverse cellular pathways including cell cycle arrest, apoptosis and DNA repair (2). In response to various stress stimuli, p53 acts as a transcriptional activator that specifically binds consensus <u>r</u>esponse <u>e</u>lements (REs; two ten base-pair half sites, RRRCWWGYYY) within its target genes to directly stimulate gene expression (2). p53 utilizes its ability to non-specifically bind and slide along the DNA to expedite the search for target REs. (3). Upon recognition of its response genes, p53 facilitates transcription at least in part by targeting the TFIID-mediated transcription machinery to the promoter (4–7). TFIID is composed of TBP and 14 <u>T</u>BP-<u>a</u>ssociated <u>f</u>actors (TAFs). TFIID recognizes and binds multiple core promoter elements (e.g. the TATA box, Initiator, and <u>d</u>ownstream core <u>p</u>romoter <u>e</u>lements [DPE]) surrounding the <u>t</u>ranscription <u>s</u>tart <u>s</u>ite (TSS) (8). Once bound, TFIID serves as a central scaffold for six basal factors, including RNA Polymerase II, to form a <u>p</u>re-<u>i</u>nitiation <u>c</u>omplex (PIC) directing transcription initation. Importantly, without the assistance of activators such as p53, TFIID’s core promoter recognition is weak and rate limiting for transcription initiation due to inefficient PIC assembly (9–16). As TFIID is responsible for transcription of at least ~90% of protein-coding genes (17), unraveling how p53 intervenes in this central step of the PIC assembly (i.e. TFIID’s binding of promoter DNA) is essential for understanding mechanisms controlling gene-specific transcription initiation throughout the cell.

Previous biochemical studies of different activators (e.g. p53 and Zta) suggested a conserved albeit poorly characterized mechanism for facilitating transcription (10, 18, 19). It has been shown that p53 binds to its REs and enhances TFIID promoter recruitment via direct contact with TBP and select TAFs (19, 20). However, p53 target genes have diverse spatial arrangements of core promoter elements and p53 REs. Each target promoter also exhibits varied affinities for p53 and TFIID (1). Therefore, it is unclear how distinct combinations of p53 REs and core promoter elements affect the dynamics and the global architecture of TFIID assemblies on target genes. Additional evidence has reported that p53 and other activators introduce structural changes within TFIID when associated with TFIIA and DNA (10, 18, 19). This activator-induced isomerization leads to enhanced TFIID binding to core promoter elements downstream of the TSS (10, 18, 19). However, the structural mechanism of p53-mediated isomerization of TFIID and its impact on PIC assembly is currently unknown.

Single molecule-based imaging has emerged as an advanced tool to discern complex dynamic behavior of large multi-subunit protein assemblies involved in regulating gene expression. A recent study utilizing real-time single molecule fluorescence microscopy provided insight that TFIID’s interaction with a synthetic core promoter DNA tightly correlates with productive transcription (21). Another single molecule report uncovered TFIIB’s dynamic binding to TFIID at the promoter (22). Additional live cell studies also suggest that the PIC composition is extraordinarily dynamic with the loading and promoter escape of RNA Polymerase II occurring every ~ 4 to 9 seconds during bursts of transcription (23). How activators such as p53 regulate TFIID’s dynamic binding to native target DNA remains elusive.

Advanced single particle transmission <u>e</u>lectron <u>m</u>icroscopy (EM) has unraveled the three-dimensional (3D) structures of TFIID and its co-complexes. Recent EM studies unmasked a structural plasticity of TFIID, likely attributed to its promoter recognition activity (20, 24–27). In particular, human TFIID features a “horseshoe” shape containing three major lobes (A, B & C) and a well-defined central cavity (28, 29). Interestingly, we demonstrated that, in the absence of DNA, three different activators, including p53, introduce a common set of local structural changes in TFIID, such as movement of lobe A towards lobe B (20). Intriguingly, human TFIID undergoes a significant structural rearrangement involving movement of Lobe A when bound to a specific promoter DNA optimized for TFIID recognition (26, 27). It remains to be determined if p53’s facilitated movement of Lobe A results in a rearranged conformation of TFIID that contributes to stable interaction between TFIID and DNA.

To uncover the mechanistic underpinnings of how p53 facilitates TFIID assembly on promoter DNA, we exploited a combination of single molecule fluorescence microscopy, biochemistry, and single particle cryo-EM analysis. These studies illuminate dynamic interactions and static global molecular architectures of p53-induced TFIID assemblies on two representative target genes (i.e. *hdm2* and *bax*). Our real-time single molecule studies show that TFIID alone infrequently assembles onto these native p53 target promoters. TFIID/DNA binding primarily occurs on the order of seconds. p53 facilitates TFIID’s promoter recruitment by enhancing multiple TFIID contacts throughout the core promoter and RE. Notably, p53 also increases the percentage of TFIID molecules displaying long-lived DNA binding. Consistent with our single molecule studies, EM structural analysis reveals that conversion of canonical TFIID to a rearranged DNA binding conformation is enhanced in the presence of p53 and DNA. TFIID/DNA binding induces p53 dissociation, effectively “liberating” the RE to allow p53-mediated recruitment of additional factors involved in PIC formation. Therefore, proper response to divergent stress signals involves conserved structural mechanisms relating to p53’s ability to dynamically stimulate TFIID’s interaction with various target promoters.

## Results

### TFIID association with native target genes in the presence of p53

We sought to further understand the molecular mechanism pertaining to activator-mediated enhancement of TFIID’s ability to bind DNA. Thus, dynamic interactions between p53, TFIID and promoter DNA were characterized via a single molecule co-localization assay using <u>T</u>otal <u>I</u>nternal <u>R</u>eflection <u>F</u>luorescence Microscopy (TIRF) (Figure 1A)(21, 22). Over the first 15’ of binding, Quantum dot-labeled TFIID (QD-TFIID) associated with only 1.5% of the Cy3/Cy5-labeled native wild-type *hdm2* promoter DNA in the absence of p53 (Figure 1B and Supplemental Figure S1, left panel), revealing a low intrinsic affinity of TFIID for DNA. Increasing the time between addition of TFIID to the DNA and the imaging of binding did not improve co-localization efficiencies, suggesting that reactions are at equilibrium (data not shown). This finding indicates that, at low concentrations (~1nM), TFIID inefficiently recognizes DNA. On the other hand, increased binding of TFIID to DNA was observed in our bulk biochemical assembly assays when much higher concentrations (100-600nM) were used as in Supplemental Figure S6B.

**Figure 1.**
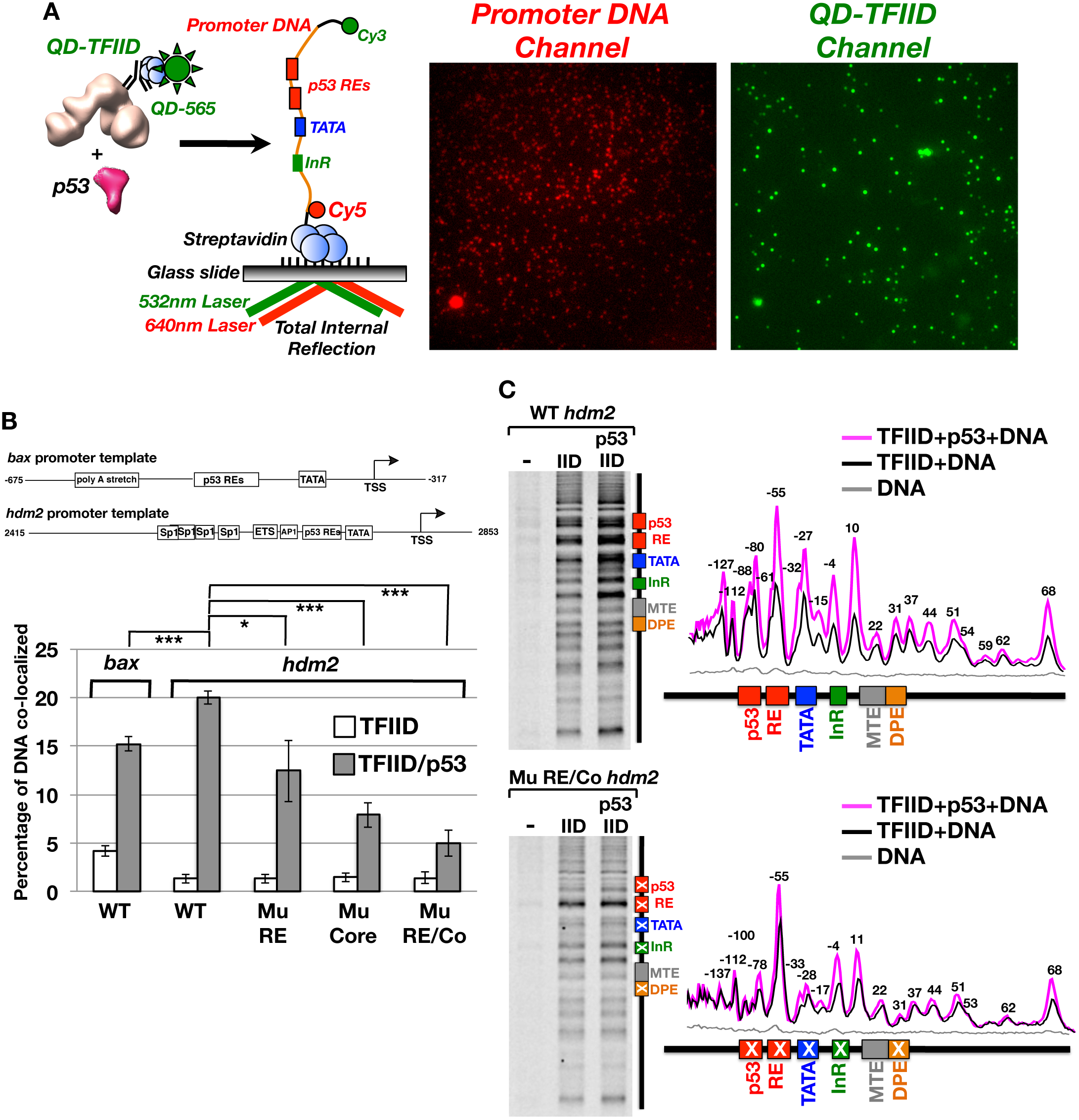
Single Molecule <u>T</u>otal <u>I</u>nternal <u>R</u>eflection <u>F</u>luorescence (TIRF) Microscopy analysis of Quantum dot-labeled TFIID binding of promoter DNA in vitro. (A) Schematic representation of an in vitro single molecule TIRF assay to monitor binding of quantum dot (QD)-labeled TFIID to fluorescently labeled *bax* or *hdm2* promoter DNA in real-time. (B) Dependence of TFIID/promoter DNA co-localization on p53, the p53 RE, and core promoter elements. Promoter DNA binding is plotted as a percentage of total DNA molecules in a field of view that are co-localized with TFIID over 15’ of binding based on TFIID–promoter DNA displacement histograms. A co-localization threshold of 84 nm was set based upon our previous study (21). TFIID binding to wild-type *bax* DNA is compared with and without p53. Additional comparisons were made for TFIID binding to wild-type (WT), Mutant p53 RE (Mu RE), Mutant Core (Mu Core), or combined mutant RE and core (Mu RE/Co) *hdm2* DNA templates. (C) xlink-Exo analysis of TFIID binding to wild-type (top panel) and Mu RE/Co (bottom panel) *hdm2* DNA. Basepair annotations (+/-2bp) above individual peaks in the line traces (right panel) represent a-4bp adjustment to reflect the known inability of Exonuclease to cut immediately adjacent to protein/DNA crosslink sites (32).

Upon addition of p53, a clear enhancement of TFIID’s association with the *bax* or *hdm2* promoter DNA was observed (Figure 1B). This result is consistent with a general increase of DNA retained by TFIID in the presence of p53 in our bulk biochemical assays (Supplemental Figure S6B). A control experiment detected no p53-mediated co-localization of DNA with the Quantum dot-labeled TAF1 antibody (0.2 ± 0.07% co-localization) in the absence of TFIID (data not shown). Therefore p53’s significant enhancement TFIID/DNA association is likely due to the direct interaction between p53 and TFIID.

To directly address the role of the p53 RE and core promoter elements in p53-mediated TFIID/DNA binding, three mutant *hdm2* DNA templates were analyzed with the same real-time co-localization assay (Figure 1B). The first mutant *hdm2* DNA template (i.e. Mu RE) harbors mutations in the response elements known to impair p53’s sequence-specific interactions (30). A mild but significant decrease of TFIID’s association with DNA was observed on the mutant p53 RE template (Figure 1B). This result suggests that enhancement of TFIID/DNA binding can occur via a mechanism that is independent of p53/RE contacts.

To assess the impact of the core promoter in p53-mediated TFIID/DNA binding, mutations (i.e. Mutant Core) known to weaken TFIID binding and inhibit transcription (22, 31) were exploited. Upon addition of p53, TFIID’s association with the mutant core template was further reduced compared to the wild-type promoter (Figure 1B). As expected, when both the p53 RE and core promoter elements were altered (i.e. Mu RE/Co), TFIID/DNA co-localization was further decreased (Figure 1B). These results indicate that the p53 RE and core promoter elements were both required for optimial association of TFIID with p53 target gene promoters.

### p53 regulates TFIID/DNA contacts within the core promoter and p53 response elements

As an alternative approach to study p53-mediated TFIID/DNA binding, we developed an in vitro formaldehyde-crosslinking/exonuclease mapping assay (xlink-Exo). Our strategy was adapted from a well-established strategy (ChIP-exo) to map out protein/DNA contacts of transcription factors in vivo at near base-pair resolution (32). Based on this analysis, TFIID makes extensive contacts with well-characterized core promoter elements (i.e. TATA, InR, MTE and DPE) on the wild-type *hdm2* DNA (Figure 1C, top panels). Addition of p53 led to elevated crosslinking of TFIID to the wild-type core promoter with select elements showing much larger changes (i.e. TATA, InR) than others (i.e. MTE/DPE) (top panels). Consistent with our single molecule data, p53 increases the specific binding of TFIID to the core promoter. Importantly, there was a clear reduction in p53’s ability to enhance contacts within mutated core promoter elements (Figure 1C, bottom panels and Supplemental Figure S2A). This finding is consistent with our single molecule assay showing reduced TFIID/DNA co-localization on the mutant core promoter (Figure 1B). Interestingly, p53 slightly increases TFIID’s contacts within the −4 and +11 regions on these mutant core promoters (Figure 1C, bottom panel and Supplementary Figure S2A). Thus our mutantions don’t completely eliminate TFIID/core promoter contacts and may explain why a small percentage of TFIID is still bound to the mutant promoters in our single molecule assays (Figure 1B).

In addition to these well-established contacts, our xlink-Exo assays also mapped TFIID/DNA interactions far upstream of TATA (−55 to −137) and downstream of the DPE (~ +37 to +68) (Figure 1C, top panel). This result is consistent with previous DNAse I footprinting studies reporting TFIID/DNA interactions upstream of TATA (−40 to −140) and downstream of the DPE (+32 to +68) at similar positions on different promoters (26, 33, 34). Many of these upstream and downstream TFIID contacts were enhanced when p53 was co-present on the wild-type *hdm2* promoter (Figure 1C). Moreover, some of TFIID’s upstream contacts (i.e. −55 to −88) reside within the p53 RE on the wild-type promoter even in the absence of p53 (Figure 1C, top panels). Mutations of the p53 RE led to a loss of TFIID’s contacts at −61 and −88, suggesting that TFIID recognizes specific sequences within the p53 RE (Figure 1C, bottom panels and Supplemental Figure S2B). Furthermore, a number of upstream TFIID contacts (−61, −79, and −88/−90) were also abrogated on the mutant core promoter (Supplementary Figure S2). This suggests that these novel upstream DNA contacts arise from TFIID’s interaction with the TATA, InR and DPE sequences. Therefore, TFIID may utilize additional uncharacterized elements to recognize promoters.

### p53 modulates the prolonged association of TFIID with DNA

In addition to p53’s ability to stimulate TFIID/DNA association, we sought to test if p53 could also stabilize TFIID’s binding to DNA. Therefore, the amount of time that QD-TFIID remains bound to a single promoter DNA before dissociating (i.e. residence time) with and without p53 was examined (Figure 2A). In the absence of p53, two populations of DNA-bound TFIID complexes displaying distinct residence times were detected on the wild-type *hdm2* template (Figure 2B). The majority (78%) of TFIID/DNA binding events were short-lived (average residence time of 6.6 ± 0.4 seconds) in the absence of p53 (Figure 2B, left panel). Notably, p53 significantly increases the percentage of long-lived TFIID/DNA bound complexes from 22% to 64% (Figure 2B, left versus right panel). Furthermore, p53 prolongs the residence time of both the short and long-lived TFIID/DNA binding events (Figure 2C). TFIID also displayed a similar stability profile on the *bax* promoter in the presence of p53 (Supplemental Figure S3). These results suggest that TFIID interacts with DNA via two intrinsic binding modes that can be modulated by p53.

**Figure 2.**
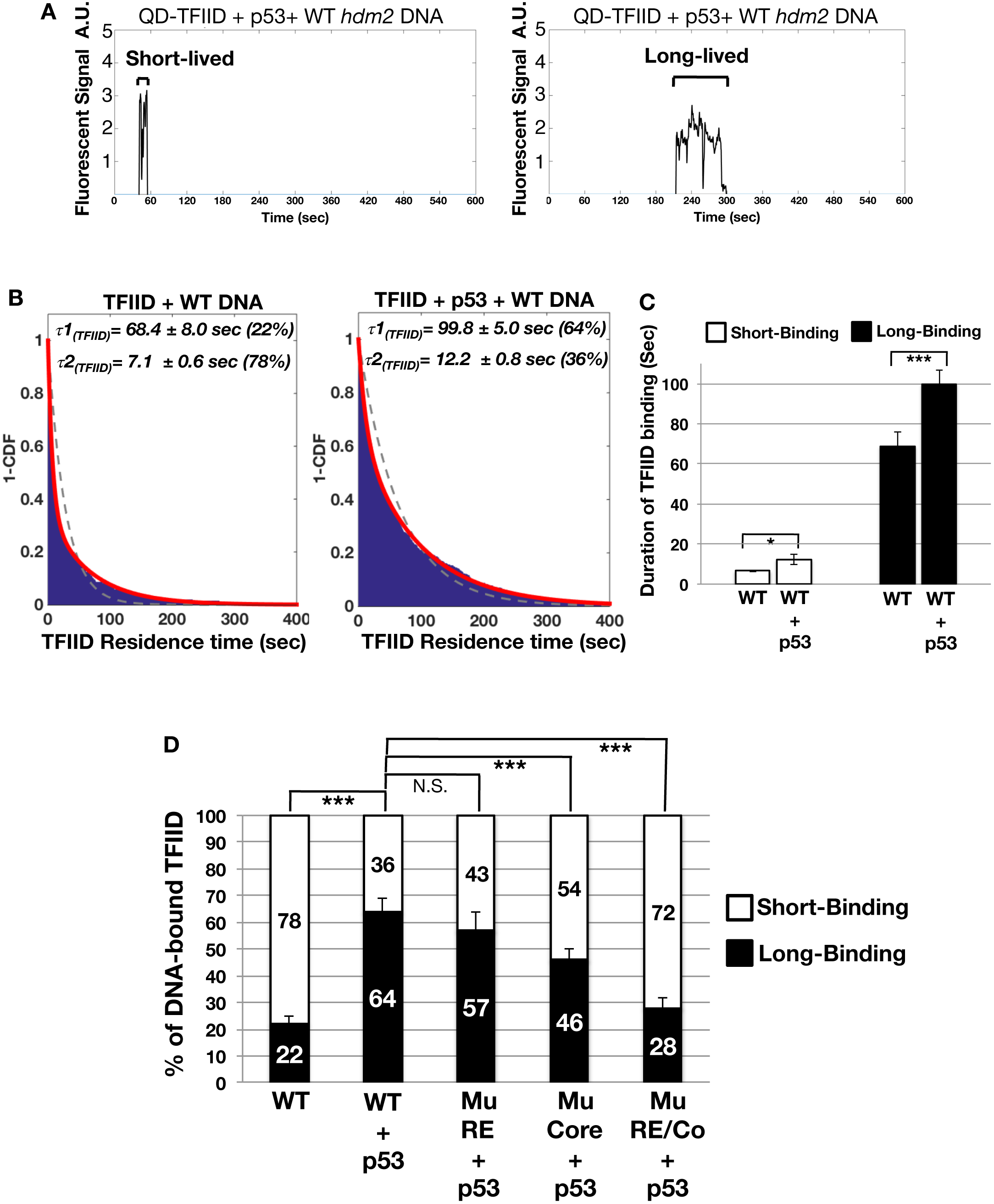
Residence time analysis of TFIID bound to promoter DNA. (A) Traces of individual co-localized TFIID/promoter DNA binding events were further analyzed to determine TFIID’s average residence time. TFIID displays a mixture of short (left panel) and long-lived (right panel) binding events. (B) 1–Cumulative Distribution Function Plots (1–CDF) of TFIID bound to DNA in the absence (left panel) and presence (right panel) of p53 were fitted to a single (gray dashed) or two-component (red solid) exponential decay model. Fitting analysis reveals that TFIID complexes bound to promoter DNA display residence times comprised of a long-(τ1)and a short-(τ2) lived component. Percentages of the long-(τ1) and short-(τ2) lived components are listed next to the residence time. (C) Bar graph comparing TFIID’s average short-and long-binding in the absence and presence of p53. (D) Percentage of short and long-lived TFIID binding events on the wild-type (RE), mutant RE (Mu RE), mutant Core (Mu Core) and mutant RE/Core (Mu RE/Co) *hdm2* promoter DNA. * = p-value < 0.05, *** = p-value <0.001, N.S. = Not significant.

We next defined how the RE and core promoter elements might regulate the relative proportion of these two TFIID/DNA binding populations in the presence of p53. Disruption of the p53 binding site on the *hdm2* promoter (Mu RE) had little effect on the distribution between these two populations (Figure 2D). This indicates that a p53-mediated increase in the percentage of TFIID molecules displaying long-lived DNA binding does not solely require contacts with the RE. In contrast, mutation of the core promoter region (Mu Core) reduced the percentage of TFIID molecules in the long-lived DNA binding population (Figure 2D). Alteration of the p53 binding site and the core promoter elements (Mu RE/Co) further lowered the percentage of the TFIID’s long-lived DNA binding population (Figure 2D). These data suggest that the TFIID’s long-lived DNA binding mode was related primarily to recognition of the core promoter with p53/RE interactions providing additional stability post-recruitment to the DNA.

Our results also reveal that p53 RE and core promoter mutations also impacted TFIID’s residence time on DNA. Mutation of the p53 RE (Mu RE) predominantly decreased the residence time of TFIID’s long-lived DNA binding without significantly affecting the short-lived events (Supplementary Figure S4B). Therefore, interactions between p53 and its RE are required for maximal stability of TFIID/DNA contacts on native target promoters. On the mutant core promoter (Mu Core) residence times for both the long-and short-lived TFIID/DNA binding events were also significantly reduced (Supplementary Figure S4). Thus, both DNA binding modes were related to core promoter recognition. Mutation of both the p53 RE and core promoter were changed (Mu RE/Co) had little effect of TFIID’s residence time on DNA (Supplementary Figure S4B). These results suggest that mutation of both the RE and core promoter elements primarily effects TFIID’s distribution between short and long-lived DNA binding modes (Figure 2D). Together with the single molecule data, our xlink-Exo studies suggest that TFIID’s remaining minimal contacts on our mutant promoter are sufficient to anchor short-lived TFIID binding to DNA.

### p53/target promoter interactions in the presence of TFIID

Our previous report documented that a TFIID variant could cooperatively increase the binding of the activator c-Jun to its binding site on promoter DNA (24). To quantitatively assess if this was the case for p53, the dissociation dynamics of fluorescently labeled-p53/DNA interactions with or without TFIID were examined (Figure 3). In the absence of TFIID, p53 primarily displayed two populations of distinct binding events on the wild-type *hdm2* template (i.e. long-lived [avg = 33.1 ± 3.9”] and short-lived [avg = 7.4 ± 0.7”], Figure 3A, left panel). Since earlier reports showed that p53 can bind DNA non-specifically (35, 36), we hypothesized that these short-lived p53/DNA binding events were non-specific interactions. To test our hypothesis, the same experiments were performed using the mutant p53 RE template (right panel). Indeed, disruption of p53’s sequence-specific binding sties on DNA led to a single short-lived population (avg = 6.8 ± 0.7”). Based upon this result, the long-lived p53/DNA binding events most likely represent sequence-specific interactions (left panel), whereas the short-lived p53/DNA binding events are non-specific in nature. Alternatively, these short-lived events could represent interactions with an unknown lower affinity p53 binding site in the promoter or binding of dimeric p53 to concensus half-sites within the mutant RE. It is of note that photobleaching of the fluorescently labeled p53 did not affect our results, since similar residence times were observed with multiple dyes of differing photostability (data not shown). In addition, a previous study detected similar short residence times of ~10-12” for p53 bound to a consensus site (37). Collectively, these findings indicate that p53 dynamically interacts with DNA via both specific and non-specific interactions.

**Figure 3.**
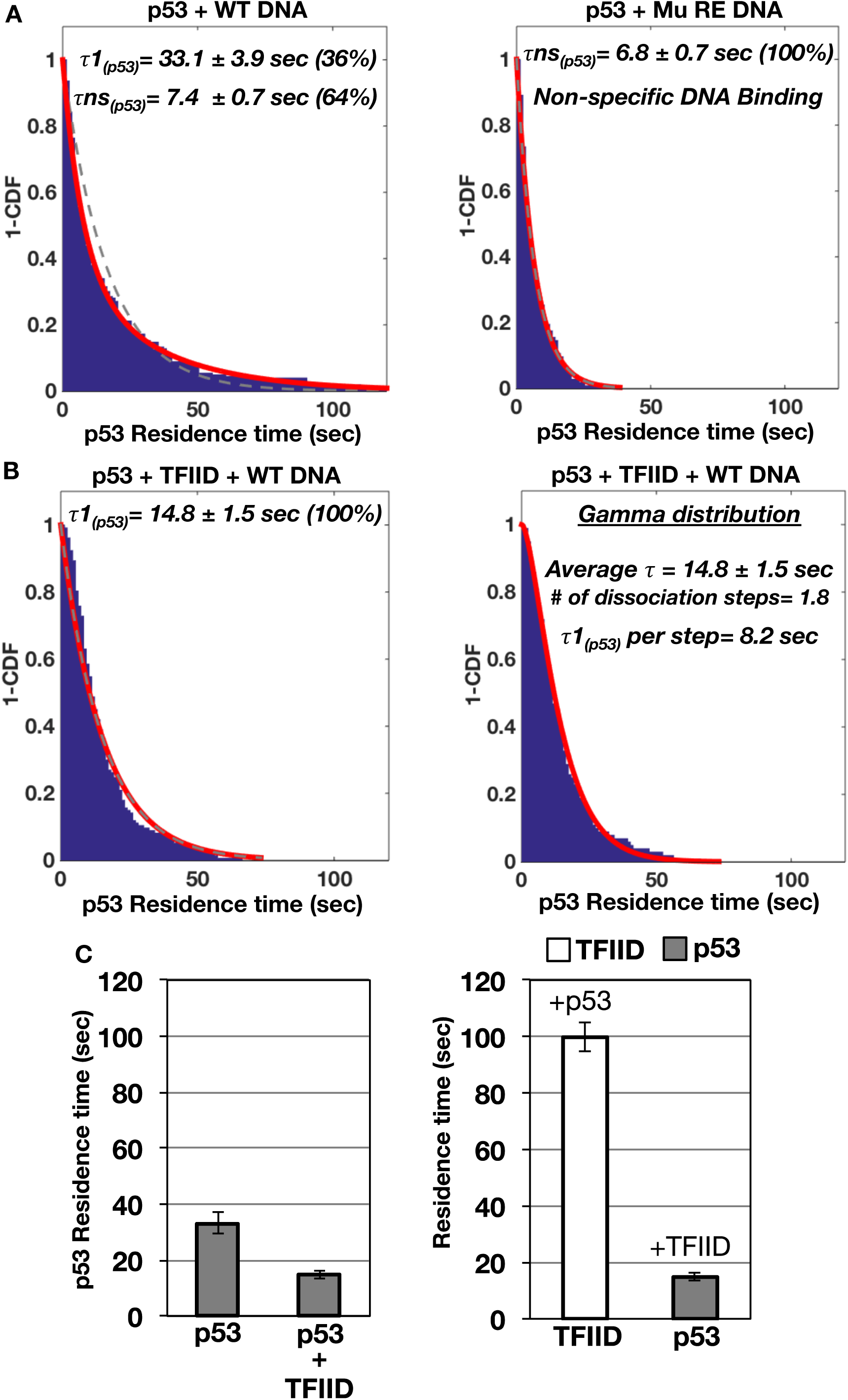
Residence time analysis of p53 bound to promoter DNA with and without TFIID. (A) Individual co-localized p53/promoter DNA binding events were further analyzed to determine the average residence time for p53 bound to wild-type (left panel) and Mutant p53 RE (right panel) *hdm2* promoter DNA. 1–Cumulative Distribution Function Plots (1–CDF) of p53 bound to promoter DNA were fit to a single (gray dashed) or two (red solid)-component exponential decay model. Based on comparisons of the *hdm2* promoter DNAs containing wild-type and mutant p53 REs, τ1 represents specific binding of p53 to the wild-type *hdm2* DNA while τns equals non-specific interactions between p53 and promoter DNA. (B) 1–CDF plots of p53 bound to the wild-type *hdm2* DNA template in the presence of TFIID were fit to a single (gray dashed) and two (red solid)-component exponential decay (left panel) or gamma distribution (right panel) model. (C) Bar graph of p53’s residence time in the absence and presence of TFIID bound to the wild-type *hdm2* DNA (left panel). Residence time analysis of TFIID and p53 bound to the wild-type *hdm2* DNA (right panel).

Importantly, addition of TFIID resulted in a single population of p53 bound to wild-type DNA with an “intermediate-lived” residence time distinct from the two populations observed with p53 alone (Figure 3B). Furthermore, we noticed that fitting of the data showed slight deviations from an exponential decay model. A previous study indicated that the existence of complex multi-step dissociation events could be discerned using a gamma distribution model to fit single molecule data (38). Indeed, we found that a gamma distribution model better represents the data (Figure 3B, right panel). This analysis shows that in the presence of TFIID, p53 dissociates from the DNA in ~2 steps. Each step contains roughly equivalent residence times of 8.2 seconds per step (average total time = 14.8 ± 1.5”). Moreover, single step dissociation kinetics was observed for p53 alone based upon gamma distribution fitting (data not shown). Therefore, TFIID alters both the rate and number of kinetic steps involved in p53’s dissociation from promoter DNA.

Mutation of the p53 RE further reduces p53’s residence time on DNA in the presence of TFIID (Supplemental Figure S5A, left panel versus Figure 3B, right panel). Disruption of TFIID’s interaction with the TATA, InR and DPE elements on the mutant p53 RE/Co DNA template does not lead to a further decrease in p53’s residence time (Supplemental Figure S5A versus S5B). Fitting of the dissociation data on the mutant templates to a gamma distribution yielded poor fits, suggesting single step dissociation kinetics for p53 on these mutant promoters (Supplemental Figures S5A and S5B, right panels).

Overall, these data suggest that TFIID binding to DNA can influence p53’s stability at the RE. TFIID’s interaction with p53 likley promotes the dissociation of p53 bound to specific sequences in the promoter (Figure 3C, left panel). Importantly, the average time of p53 bound to the wild-type *hdm2* template (~15”) is significantly shorter than TFIID’s long-lived DNA binding time (~100”) (Figure 3C, right panel). These dynamic studies strongly suggest that p53 transiently escorts TFIID to the promoter DNA. Importantly, p53 is not required to remain bound for TFIID’s maximum stability on promoter DNA.

### The conformational status of TFIID when co-present with p53 and promoter DNA

A recent cryo-EM study discovered that human TFIID switches from a canonical to a rearranged conformation upon binding to the <u>S</u>uper <u>C</u>ore <u>P</u>romoter (SCP), a DNA fragment optimized for interaction with TFIID (26, 31). Thus far our single molecule data indicates that p53 stimulates binding of TFIID to DNA. Therefore, we hypothesized that TFIID could form a rearranged DNA binding conformation with p53 and promoter DNA. Therefore, we set out to determine TFIID’s global structural architecture when p53 and target promoter DNA were present. As the first step, unbiased reference-free 2D classification experiments were performed to examine conformational states of TFIID in the presence of p53 or p53*/bax* DNA (Figure 4A). The movement of Lobe A from Lobe C to Lobe B is the hallmark of the rearranged DNA binding conformation of TFIID (26). Therefore, three conformational states of TFIID represented by corresponding views of 2D class averages were defined based upon the position of Lobe A relative to Lobes C and B (top panels, i.e. canonical, transition, and rearranged state). We found that addition of p53 led to an elevated percentage of particles within the corresponding view displaying the rearranged TFIID form compared to TFIID alone (Figure 4A, bottom panel). This finding implies that the formation of TFIID’s rearranged state is affected by p53. This result is consistent with conformational changes observed in our previous negative stained structure of a TFIID/p53 co-complex (20). Furthermore, the 2D classification analysis showed that the populations of both the transition and rearranged forms increased when p53 and DNA was added (bottom panel). Therefore, these findings suggest that TFIID’s conformational switches can be regulated via target gene promoters and/or its interaction with p53.

**Figure 4.**
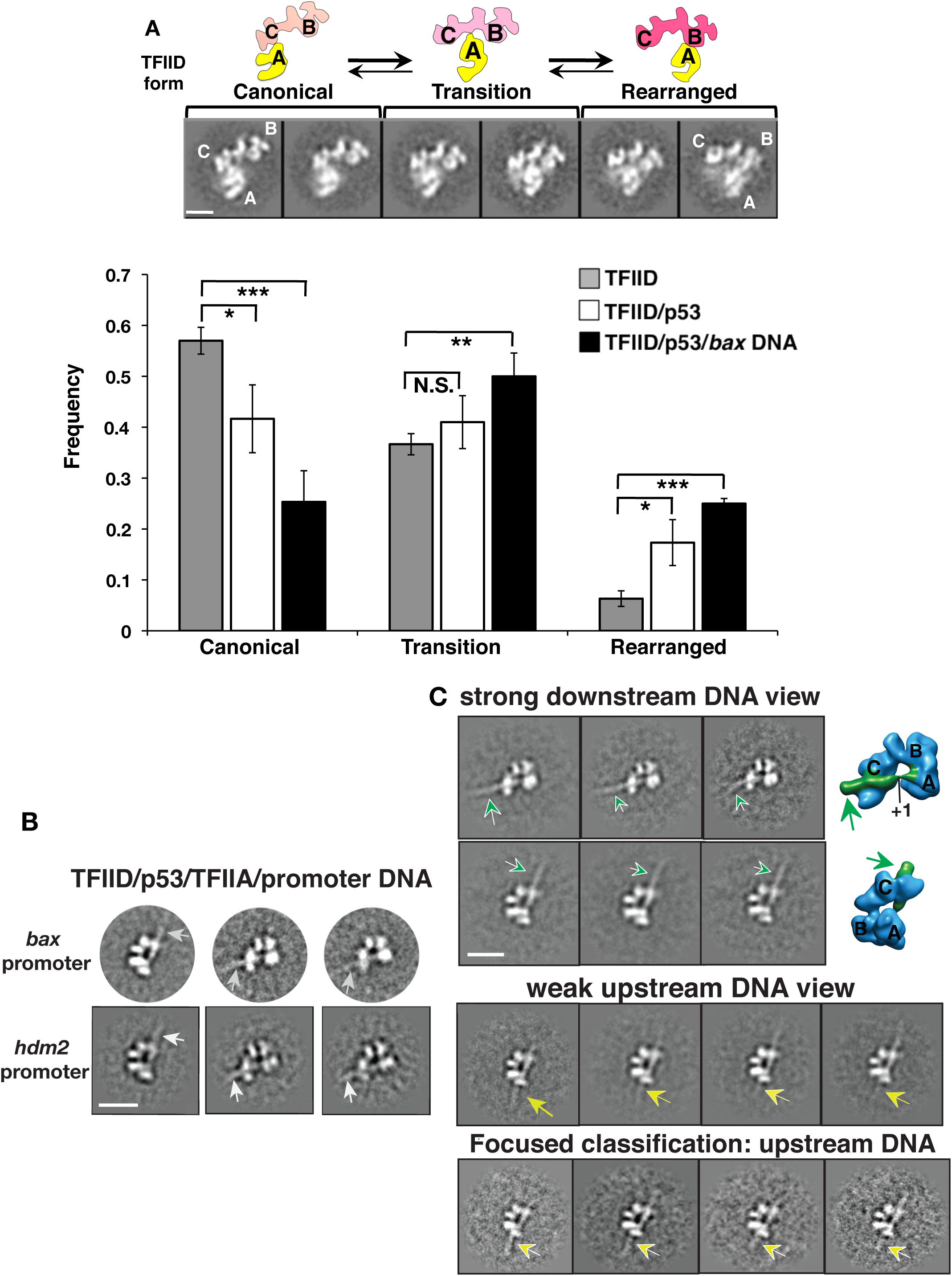
Structural EM analysis on p53’s impact on TFIID bound to native target gene promoters. (A) TFIID’s structural plasticity as defined by the position of its Lobe A (26) (top panels, the major lobes of TFIID are labeled as A, B, and C). TFIID’s conformations are classified into three groups: Canonical, Transition and Rearranged state as shown. Reference-free 2D classification analysis on negative stain EM datasets showing populations of different TFIID conformations in the absence of p53 (gray), presence of p53 (white), and with p53 +*bax* DNA (black). The scale bar represents 100Å. (B) Three representative reference-free 2D class averages of p53/TFIID/TFIIA bound to either the *bax* or *hdm2* DNA are shown based upon cryo-EM particle datasets. The scale bar is 200Å. (C) Distinct 2D class averages showing views of the assemblies bound to the *bax* promoter DNA fragment downstream (top panel, DNA highlighted in green arrows) and upstream (middle panel, DNA highlighted in yellow arrows) of the transcription start site (TSS, +1) are presented respectively. The orientation of the core promoter elements bound by TFIID were previously mapped in the 3D structure of TFIID/TFIIA bound to the <u>s</u>uper-<u>c</u>ore <u>p</u>romoter (SCP) DNA (26). The corresponding views of the 3D structure of TFIID/TFIIA/SCP DNA are shown (far right) to indicate the DNA orientation. The strong 3D density highlighted in green represents the promoter DNA comprising the initiator, TSS and downstream promoter elements. Since the upstream promoter DNA was weakly visible, focused classification analysis (bottom panel) were performed using those corresponding class averages (middle panel) to clarify the presence of the *bax* promoter DNA upstream of the TATA box (highlighted by yellow arrow). The scale bar is 200Å.

A previous biochemical study reported that p53 stimulated the assembly of TFIIA and TFIID on a synthetic promoter DNA (19). Interestingly, unlike p53, TFIIA alone promotes the canonical form of TFIID (26). Thus, to assess the structural framework of TFIID in the presence of p53 and TFIIA, two TFIID/p53/TFIIA assemblies on the native *hdm2* and *bax* promoter fragments (488 bp and 500 bp) were obtained (Supplemental Figure S6). To visualize both conformations and the DNA, cryo-EM data of these assemblies were obtained for reference-free 2D classification analysis. Some class averages showing the presence of promoter DNA spanning the central cavity of TFIID were observed (Figure 4B). Moreover, both assemblies displayed a similar rearranged TFIID architecture, despite the different organization of their promoters. Each template harbors non-consensus core promoter elements with different spatial arrangements between TATA and the Initiator (Figure 1B, top panel) (8, 39, 40). Combined with our 2D classification analysis (Figure 4A), these structural studies indicate that TFIID binds DNA in a common rearranged conformation in the presence of p53 and TFIIA.

Next, to identify the relative DNA orientation and any distinct structural features, we compared the class averages of our two assemblies with the previous structural analysis of TFIID/TFIIA/SCP DNA containing a mapped DNA orientation (26). Our co-complexes displayed clear views of TFIID stably bound to the *bax* DNA downstream of the TSS (Figure 4C, top two rows). In contrast, upstream promoter DNA was weakly visible in our averages (middle panel). These results suggest that the p53 RE upstream of the TATA box element is flexible within the co-complex. To define promoter DNA upstream of TATA, focused classification on a class average displaying flexible upstream DNA (middle panel) was performed as previously described (26). This analysis revealed a clear density for the upstream promoter DNA (highlighted by yellow arrow, bottom panel). Given the strong density for downstream DNA within the assembly, these results imply that TFIID serves as a strong anchor for downstream promoter DNA during PIC assembly.

The 2D class averages of TFIID/DNA/p53/TFIIA assemblies exhibit a similar topology to the averages from TFIID/TFIIA bound to the SCP DNA (refer to Figure 3 in (26)) raising two major points. First, TFIID adapts a common DNA binding conformation to bind various p53 target gene promoters. Second, structural rearrangements within TFIID related to DNA binding may cause p53 to dissociate, as supported by our single molecule dynamics results (Figure 3). To test these hypotheses, single particle 3D reconstruction was conducted to determine the 3D structure of the TFIID/TFIIA/p53/*bax* DNA co-complex (Supplemental Figure S7A). The overall structural architecture of our co-complex is comparable to the 3D structure of TFIID/TFIIA/SCP DNA. Our 3D reconstruction was also verified by analyzing its 3D projections with their matching reference-free 2D averages (Supplemental Figure S7B). The same observation was also obtained with the 3D structure of the TFIID/TFIIA/p53/*hdm2* promoter DNA (Supplemental Figure S8). These results suggest that TFIID utilizes one common rearranged form to bind various promoters.

Our 3D reconstruction did not reveal any prominent extra densities that would most likely represent the p53 protein. This indeed suggests that p53 dissociates from the TFIID/TFIIA/*bax* DNA co-complex. Alternatively, p53 could have a transient interaction with the assembly, which led to the EM density being averaged out during analysis. Since we previously determined the 3D structure of the p53 bound TFIID co-complex without DNA (20), we attempted to capture the p53-bound state of TFIID in the cryo-EM dataset of our TFIID/TFIIA/p53/*bax* DNA assembly. Thus, 3D reconstructions were performed using the structure of p53/TFIID as an initial reference volume. A poor resolution 3D reconstruction of p53/TFIID was obtained, indicating a small subset of the single cryo-EM particles (Supplemental Figure S9). This observation suggests that the majority of TFIID complexes have switched to the p53-free, DNA-bound state when p53 and DNA were co-present. Collectively, these structural studies support the observations from our single molecule analyses, illustrating that p53 dissociates from TFIID and DNA when recruited to target promoters. Our work also implies that when p53 stimulates TFIID binding to DNA, a common rearranged DNA binding scaffold is formed on various target genes.

## Discussion

### TFIID infrequently interacts with native p53 target genes in the absence of additional factors

Our single molecule assays reveal that TFIID poorly associates with native p53 target promoters in the absence of p53 (Figure 1B). TFIID’s inefficient DNA binding activity may be one way to prevent spurious basal expression of these stress response genes. TFIID/promoter interactions are likely repressed in part due to the TAF1 N-terminal domain (TAND) which occupies TBP’s concave DNA binding surface (41–44). This TAF1/TBP interaction is also thought to control the TFIID’s global conformation to regulate promoter interactions. TFIID’s canonical conformation is non-permissive to DNA binding (26, 27) (Figure 5 top panel, left side). Researchers postulate that the TAF1 TAND acts to tether TBP to TFIID’s lobe C in the canonical form (27). Large-scale structural rearrangement of TFIID dramtically shifts TBP’s positon within TFIID to allow simultaneous contact of TBP with the upstream TATA and TAF1 with downstream core promoter elements (Figure 5 top panel, right side). Thus TAF1/TBP interactions repress the large-scale structural rearrangement within TFIID required for TBP to bind the TATA element in a core promoter.

**Figure 5.**
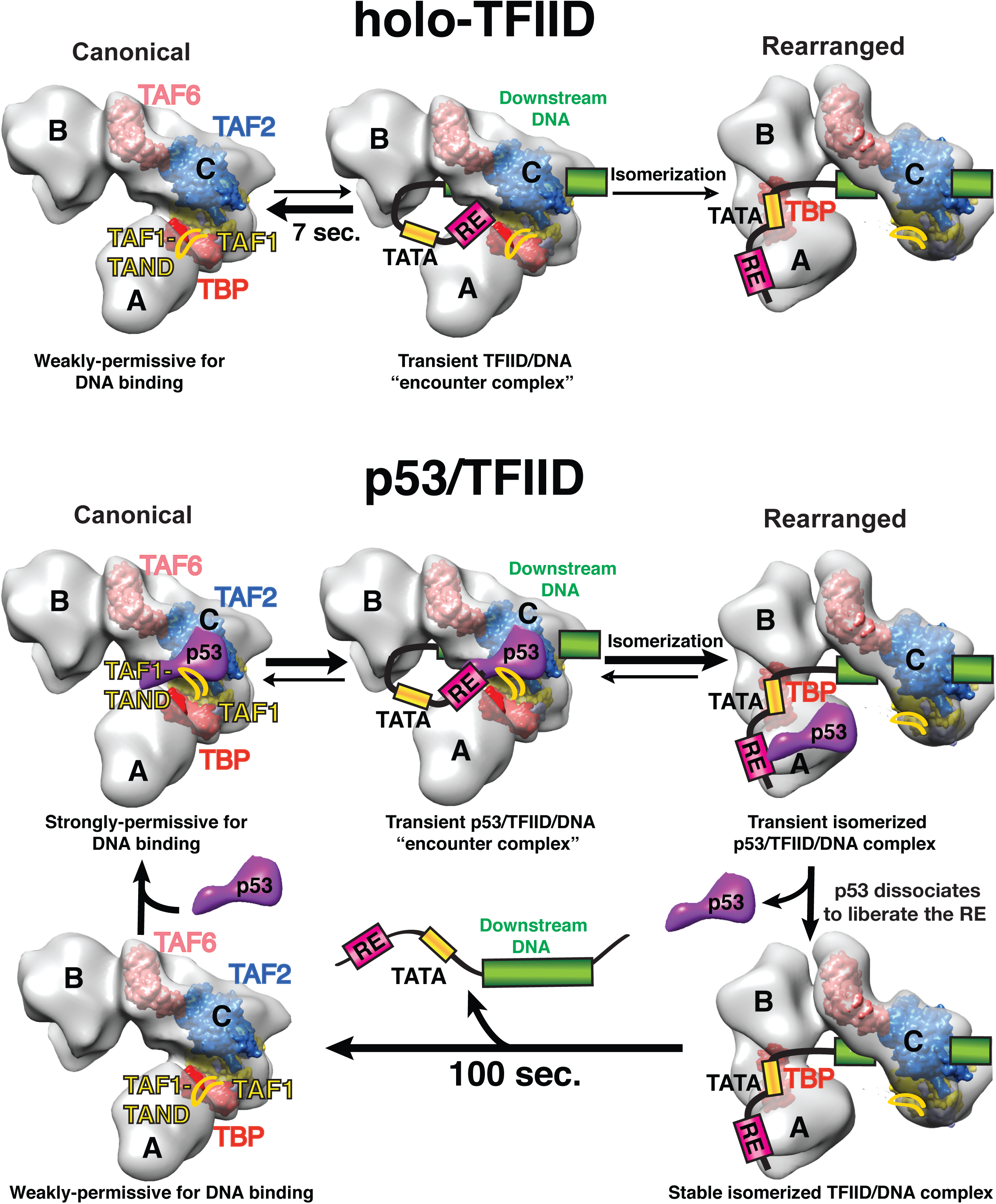
A representative model of how TFIID engages the core promoter at various p53-responsive promoters. A proposed model of how p53 aids TFIID in core promoter recognition is presented based upon our current and previous studies (20). In the absence of p53, TFIID primarily exists in the canonical conformation which is weakly permissive for DNA binding due in part to interactions between the TAF1 TAND and TBP. The canonical form of TFIID can infrequently and transiently interact with DNA. In rare occurances, canonical TFIID isomerizes to a structurally rearranged form that involves severance of TAF1 TAND/TBP contacts and movement of TBP from lobe A/C to lobe B. In the presence of p53, the TBP/TAF1 TAND interaction is disrupted leading to a TFIID complex that is highly permissive to isomerization and stable DNA binding. Once TFIID structurally rearranges and binds DNA via movement of TBP from lobe A/C to lobe B, p53 disengages TFIID through loss of its contacts with Lobe A and C. Dissociation of p53 from the TFIID/promoter DNA scaffold results in “recycling” of the p53 RE for further assembly of the PIC components.

TFIID’s infrequent association with p53 target gene promoters occurs via distinct short and long-lived DNA binding modes (Figures 2B, 2C and Supplementary Figure S4B). The majority (78%) of these TFIID/DNA binding events last just a few seconds (~7”) (Figures 1B, 2B and 2D). This short-lived DNA binding population may represent a transient non-productive TFIID encounter complex in the canonical form where TAF1 prevents stable TBP/TATA interactions (Figure 5). In such a scenario, TAF contacts with the initiator and DPE may be sufficient to temporarily anchor TFIID to DNA. Due to this transient dynamic interaction behavior, we were likely unable to obtain the 3D structure of those short-lived DNA-bound TFIID complexes via cryo-EM.

### DNA binding and structural features of TFIID are regulated by p53

A number of factors, including activators (i.e. c-Jun and VP16) and TFIIA, can stimulate TFIID/promoter binding by de-repressing the TAF1-mediated inhibition of TBP (19, 45–48). TFIIA has also been found to aid conversion of canonical TFIID to the rearranged DNA binding form likely via disruption of TBP/TAF1 TAND interactions (26, 27). It was previously unclear if activators could also promote structural rearrangement of TFIID to stimulate promoter association. Our past work has shown that p53 binds stably to TFIID and targets TAF1 in the absence of DNA (20). We also found that p53’s interaction with TFIID also results in movement of Lobe A towards Lobe B in a manner analogous to a structurally rearranged DNA binding form of TFIID (20). Therefore we postulated that DNA binding may be increased in this structurally rearranged pre-assembled p53/TFIID complex. Indeed our single molecule data along with previous bulk biochemical experiments indicate that p53 can dramatically stimulate the binding of TFIID to DNA (Figure 1B and (19)). We also find that conversion of canonical TFIID to the structurally rearranged form is enhanced in the presence of p53 and DNA (Figure 4A). Thus we speculate that p53, whose acidic activation domain is highly homologous to VP16 (49), may potentially disrupt TAF1/TBP interactions to aid in TFIID structural rearrangement and DNA binding (Figure 5, bottom panel).

### p53 promotes isomerization of TFIID to stimulate long-lived binding to DNA

Cryo-EM of a TFIID/TFIIA/DNA complex has revealed a number of TBP/TAF contacts with core promoter elements spanning from ~ −33 to +32bp surrounding the transcription start site (27). Many of these contacts, especially TBP/TATA, are only possible due to TFIID’s structural rearrangement or isomerization upon DNA binding. Therefore we anticipate that the p53-mediated binding of TFIID to DNA is also likely a multi-step isomerization process involving contacts with multiple core promoter elements (Figure 5, bottom panel). Indeed our x-link Exo analysis revealed numerous TFIID crosslinks throughout the promoter that are enhanced by p53 (Figure 1C). Many of these *TFIID/hdm2* DNA contacts were strictly conserved in the TFIID/TFIIA/super core promoter DNA cryo-EM structure (27). Consistently, the architectures of TFIID bound to the *hdm2* and supercore promoters are equivalent (Supplemental Figures S7 and S8). This suggests that TFIID in the presence of p53 can specifically bind our native *hdm2* promoter to adopt a common DNA binding conformation.

In the presence of p53, TFIID associates with DNA robustly. Analysis of single molecule dissociation kinetics also revealed two TFIID/core promoter DNA binding modes with interactions predominantly being long-lived (~100”) (Figures 2B and 2D). Therefore, p53 shifts the TFIID population from short to long-lived DNA binding, likely acting to facilitate TFIID’s known isomerization on promoters (Figure 5, bottom panel). This isomerization step may possibly permit stable interaction of TFIID with the upstream DNA, TATA box and downstream DNA resulting in long-lived DNA binding (Figure 5, bottom panel). We speculate that the p53- mediated long-lived DNA-bound TFIID complex is structurally related to the stable TFIID/TFIIA/promoter DNA complex depicted in our cryo-EM studies. In support of this model, previous biochemical assays have shown that activators induce conformational changes within TFIID that enhance multiple contacts with promoter sequences downstream of the TSS (10).

Mutation of the p53 RE only affected the residence time of TFIID’s long-lived DNA binding population (Supplementary Figure S3). Importantly, the percentage of long-lived DNA binding events was essentially unchanged on the mutant RE template (Figure 2D). This suggests that isomerization of TFIID does not depend on p53 contacts with the RE. Given that distinct TFIID contacts surrounding the p53 RE were identified via our xlink-Exo assays (Figure 1C), we speculate that this upstream DNA is involved in both binding p53 and stabilizing interaction of an isomerized form of TFIID with the promoter. These upstream contacts may be possibly mediated by the TAF4 subunit of TFIID, which was previously shown to bind ~70 bp of DNA with a weak sequence preference (50).

### TFIID isomerization upon DNA binding liberates p53 from the RE

Our single molecule data indicate that TFIID reduces p53’s residence time on DNA (Figures 3A and 3B). This result is consistent with previous DNAse I footprinting studies showing decreased binding of p53 to REs in the presence of TFIID (19). More importantly, p53’s residence time on DNA is 6.7 fold shorter (~15”) than TFIID’s (~100”) (Figure 3C, right panel). Therefore p53 disengages the promoter before TFIID during long-lived DNA binding events. In our EM structures, we are also unable to detect density associated with p53 in the rearranged DNA bound TFIID further confirming our single molecule findings (Supplemental Figures S7, S8 and S9). Previous structural analysis shows that p53 bridges lobes A and C of TFIID in the absence of DNA (20). Thus, we speculate lobe A’s movement away from lobe C upon DNA binding disrupts p53/TFIID contacts inducing p53’s dissociation from the assembly (Figure 5, bottom panel).

While p53 dissociates from the TFIID/DNA scaffold, it is highly likely that p53 will rapidly re-associate with the complex, particularly at the high concentrations (600nM) used in Supplementary Figure S6. We suspect that p53’s rapid removal could consequently lead to a faster “repurposing” of the RE for additional rounds of p53-mediated recruitment of other basal transcription factors such as TFIIB (51, 52), RNA Pol II (53), and TFIIH (49, 54, 55). In this scenario, stable prolonged binding of p53 to a TFIID/DNA complex would negatively regulate PIC formation, because p53 would occupy the RE and inhibit subsequent recruitment of a p53/basal factor co-complex.

### TFIID/promoter binding dynamics in relation to transcriptional burst kinetics

Once p53 loads TFIID onto our native p53 target gene promoters, TFIID remains bound for only approximately 100 seconds likely due to interactions with suboptimal or non-consensus core promoter elements (Figure 2). Notably, our residence times for TFIID at the promoter were measured in the absence of additional PIC components. Most likely, additional factors such as TFIIA, TFIIF, and RNA Polymerase II will alter the stability of TFIID and p53 as the PIC forms on promoter DNA. However, recent in vivo work suggests that transcription complex formation is highly dynamic. Consistent with our data, yeast TBP occupies the HSC82 heat shock promoter for 60 seconds in vivo (56). In human tissue culture cells, RNA Polymerase II loads onto promoters every ~4 to 9 seconds to form convoys on an active gene (23). These bursts of RNA Polymerase II convoys last for approximately 100 seconds before the promoter becomes inactive again (23). Therefore it is tempting to speculate that the long-lived TFIID residency (~100 seconds) on our native target promoters could be related to RNA Polymerase II convoy formation (~100 seconds) and transcriptional bursting. TFIID’s dynamic binding to promoters could be a mechanism to allow rapid shut down of stress response gene expression after re-establishment of cellular homeostasis.

TFIID can also direct infrequent loading of RNA Polymerase II convoys on mutant TATA containing promoters leading to bursts of transcription (23). Interestingly, the duration of RNA Polymerase II convoys on the mutant TATA promoter is only slightly less than that on a wild-type TATA template (~60-80 seconds) (23). Our study also revealed infrequent p53 mediated binding of TFIID on our mutant core promoter DNA lasting for a significantly long time (~64 seconds) (Figure 2D and Supplementary Figure S3). Therefore we suspect that TFIID stably loaded onto our mutant core DNA is fully competent to direct low levels of infrequent transcription. Overall, our studies indicate that p53, and potentially activators in general, serve as escorts to dynamically recruit and load the basal transcription machinery onto DNA via complex interactions between the RE, the core promoter and bound basal factors (i.e. TFIID). Our work also reveals how p53 stimulates TFIID binding to different promoters that comprise variable arrangements of p53 binding sites and core promoter elements. More importantly, our combined functional and structural studies are crucial for understanding how eukaryotic transcription complexes dynamically assemble on different protein-coding gene promoters.

## Materials and Methods

### Reagents and protein purification

Experimental details, including reagents, protein purification of the p53/TFIID/TFIIA/DNA co-complexes and fluorescent-labeled proteins/DNA used in this study along with our custom-built TIRF microscope setup, are described in the Supplemental Materials.

### In vitro immuno-assembly assay

HeLa cells (32 liters) were grown in suspension culture with 1X DMEM plus 5% newborn calf serum for each assay. Nuclear extract was prepared and fractionated with phosphocellulose P11 (P-Cell) resins exactly as previously described (20). P-Cell column fractions eluting at 1M KCl/HEMG buffer (pH 7.9, [20 mM Hepes, 0.2 mM EDTA, 2 mM MgCl_2_, 10% Glycerol, 1 mM DTT and 0.5 mM PMSF]) were pooled, dialyzed to 0.3 M KCl/HEMG buffer, added with final concentration of 0.1% NP40, and then immunoprecipitated overnight at 4°C with an anti-TAF4 mAb covalently conjugated to Protein G Sepharose beads (GE Healthcare Life Sciences). TAF4-immunoprecipitates were extensively washed with 0.65 M KCl/HEMG buffer, 0.3 M KCl/HEMG buffer and 0.1M KCl/HEMG buffer (containing 0.05% NP40 buffer and 10 μM Leupeptin) and split into four assembly reactions as indicated in Figure 1. Three reactions of TFIID-bound resins were incubated with 6-fold molar excess of promoter DNA. Next, two TFIID/DNA reactions were further added with 10-fold molar excess of p53 alone and/or TFIIA. Four total reactions were incubated at 4°C for 2 hours prior to elution of the co-complex from the TAF4 mAb affinity resin. TFIID/factor/DNA-bound resins were washed 5 times with 0.1 M KCl/HEMG buffer (containing 0.1% NP40 and 10 μ M Leupeptin) followed by elution with a peptide (1 mg/ml) in 0.1 M KCl/HEMG buffer (plus 0.1% NP40 and 10 μM Leupeptin) recognized by the TAF4 mAb. Eluates were analyzed with 4-12% SDS-PAGE (NuPage gel system, Thermo Fisher) and visualized by silver staining analysis.

### Xlink-Exo assay

Exonuclease assays were carried out in 10μl reactions containing a modified 50 mM KCl/HEMG buffer (pH 7.9, [12.5 mM Hepes, 0.05 mM EDTA, 6.25 mM MgCl_2_, 5% glycerol] plus 1% DTT, 0.05 M PMSF, and 0.01% NP-40), IRdye800 (IDT) antisense strand labeled *hdm2* promoter DNA (9nM), p53 (90nM, monomer), and TFIID (4nM). After incubation at room temperature for 20’, reactions were crosslinked with formaldehyde (1% final) for 5’ and stopped by addition of Tris-HCl (100mM final, pH 8.0). Next, each reaction was mixed with Exonuclease III (50 Units, NEB diluted in a buffer containing MgCl_2_ [16.25 mM final]), and incubated at 37^o^C for 1 hour. Proteinase K (10μg) was then added into each reaction and incubated at 70^o^C for 3 hours. Reactions were mixed in formamide loading dye, heated at 95^o^C and resolved on a 8% Polyacrylamide-TBE gel containing 7M urea. A series of known 5’ truncated IRdye800 (IDT) antisense strand labeled wild-type *hdm2* DNA fragments were generated and used as molecular standards to determine the size of Exonuclease digested bands seen on our mapping gels.

### TIRF Single Molecule Imaging

Passivated glass coverslips (see Supplemental materials) were assembled into a multichannel flow cell using double sided tape (3M). Flow cells were mounted on the microscope and streptavidin was added to the flow cell to coat surfaces. Each flow cell holds ~50 ul of sample. Reference beads that serve as fiducial marks were immobilized on the bottom surface of the flow cell. Biotinylated Cy3/Cy5 promoter DNA in 1X PBS was then loaded into the flow cell and bound over time specifically to the streptavidin that was non-covalently attached through biotin linkages on the top and bottom surfaces. Promoter DNA templates were briefly imaged at 3HZ to determine surface attached DNA densities that were sufficient for localization of single DNA molecules. Unbound biotinylated Cy3/Cy5 promoter DNA was washed away. Promoter DNA templates were then imaged at 3HZ. The surface attached Cy3/Cy5 promoter DNA was then photobleached. Reactivation of the Cy3/Cy5-labeled promoter DNA was not evident over the timescale of our experiments (typically 15-20 minutes). Qdot 565-labeled TFIID (1 nM) in the absence and presence of unlabeled p53 (8 nM, monomer) was then added to the flow cell in a modified 50 mM KCl/HEMG buffer (pH 7.9, [12.5 mM Hepes, 0.05 mM EDTA, 6.25 mM MgCl_2_, 5% glycerol] plus 1% DTT, 0.05 M PMSF, and 0.01% NP-40) and immediately imaged at 3Hz at 23^o^C. An oxygen scavenging system consisting of 0.4% b-D-Glucose, 1 mg/ml glucose oxidase (Sigma), and 0.4 mg/ml catalase (Sigma) was used for imaging. For experiments to measure p53’s interaction with promoter DNA, AlexaFluor 555, Cy3-or Cy5-labeled p53 (8nM, monomer) in the absence or presence of unlabeled TFIID (1 nM) was added to the flow cell containing surfaced attached mapped promoter DNA and immediately imaged at 3Hz. AlexaFluor 555, Cy3-and Cy5-labeled p53 had equivalent residence times on promoter DNA indicating that photobleaching of molecules didn’t significantly affect results.

### Single Molecule Co-localization Analysis

Fluorescent spots in each frame for the movie of the promoter DNA, TFIID or p53 were mapped using 2D Gaussian fitting methods (21, 57, 58) to determine the location (x, y) at subpixel resolution. Co-localization analysis was performed as in (21) with the following modifications. Coordinates of fluorescent spots were adjusted based upon movement of the fiducial marks (i.e. reference beads) over time. Stage movement/drift was typically less than 200 nm over the time of imaging (~20 minutes). Fluorescent spots that were located within 84 nm (1 pixel) throughout the movie were grouped together in a cluster. The positions of all spots within a cluster were averaged to obtain the overall location of an individual cluster of molecules. Position maps of the clustered surface attached DNA molecules were then compared with position maps of either the TFIID or p53 clusters. Clusters of TFIID or p53 molecules within 3 pixels (252 nm) of a DNA cluster were found. Displacements of TFIID or p53 clustered molecules with a maximum equal to (0,0) in the x and y positions are indicative of non-random co-localization between DNA and TFIID or p53 clusters. Displacements between the x,y positions of the TFIID or p53 clusters and the DNA clusters were then plotted as a 2D histogram with a bin size of 50 nm. 2D Gaussian fitting of the 2D histogram yields σ which is equivalent to the variation in the displacement of TFIID or p53 within 252 nm of a DNA molecule. A co-localization threshold was set to 1.8σ (84 nm).

### Single Molecule Residence Time Analysis of TFIID and p53 on promoter DNA

For each TFIID or p53 cluster that co-localized with a DNA cluster (i.e. within 84 nm), we the plotted the amplitude from the 2D Gaussian fit of every fluorescent spot within the TFIID or p53 cluster as a function of time. Homemade MATLAB software was then used to analyze the rise and the fall of the amplitude of the fluorescent signal to determine the appearance and corresponding disappearance of each spot over time. The residence time for each DNA binding event is defined as the difference in time between the appearance and disappearance of a TFIID or p53 fluorescent spot. The fluorescent signal from TFIID or p53 had to last for at least 1.5” to be considered as a discrete binding event in our analysis. Due to rapid stochastic blinking of the quantum dot, a dissociation event was only considered to have occurred if the quantum dot signal remained in the off state longer than 4.5”. At least 300 binding events of co-localized TFIID or 100 binding events of p53 clusters were analyzed for each particular promoter DNA construct. For a subset of molecules (~100), the data was processed by hand to determine the appearance and disappearance of fluorescent spots. Comparison of the residence times generated by our automated software and by hand yielded negligible differences. Histograms of residence times for TFIID or p53 bound to promoter DNA were fit to a two-component exponential decay model to determine the average residence time of TFIID or p53 bound to promoter DNA (58). If the difference between the fitted residence times for each individual component in the two-component model varied less than 20%, histograms were re-fitted using a single-component exponential decay model (58). For Figures 3 and S5 data was additionally fit to a gamma distribution (38) via custom MATLAB software.

### 2D Classification Analysis of TFIID co-complexes via negative stain EM

4 μL (10-20 ng in total amount) of fresh TFIID/p53/bax DNA assemblies in 0.1M KCl/HEMG buffer (20mM HEPES, 0.2 mM EDTA, 2mM MgCl_2_, 10% Glucose [final pH 7.9]) was applied directly onto a thin carbon film supported by holey carbon on a 400-mesh copper grid (Pacific Grid tech.) for 3’, which was freshly glow-discharged. After incubating the sample on the grid for 3’, the sample grid was stained with five successive 75 μL drops of 1% Uranyl Formate.

The negative stain image data were collected with a Tecnai F20 TWIN transmission electron microscope operating at 120 keV at a calibrated magnification of 62,000x with a defocus range of −0.5μm to −2.5μm using a Tietz F416 (4K × 4K) CCD camera (resulting in a 2.82 A/pixel) under the dosage 39.98 e^-^/Å. A total of 14,000 particles (for TFIID alone), 13,855 particles (for TFIID/p53), and 19262 particles (for TFIID/p53/bax DNA) were manually picked using Boxer (EMAN; (59)). These particles were aligned and classified in a reference-free fashion as previously described (24) for 8 iterations. The similar measurement of Lobe’s A position relative to the lobes B and C was employed as in (26). Based upon the position of Lobe A in relation to Lobe C v.s. Lobe B (26), three conformational groups of TFIID were defined (i.e. canonical, transition, and rearranged state) in this study. The canonical form is represented by Lobe A retaining its attachment to Lobe C. The transition state is represented by Lobe A beginning to detach and move in between Lobe C and B. The rearranged form is represented by Lobe A reattaching to Lobe B. Next, individual classums from the 5^th^, 6^th^, 7^th^ and 8^th^ iteration for each TFIID/p53 prep with or without the *bax* DNA were examined carefully to select 2D class averages displaying views of these three conformational groups. The number of 2D averages in each conformational form v.s. total averages was calculated to yield the percentage of TFIID’s three conformations in each TFIID/p53 prep with or without the *bax* DNA. Three independent protein purifications and EM data acquisition/analysis were performed to obtain the standard deviation for the figure. A two sided student’s T-test was used to determine statistical significance.

### Cryo-EM data collection and image analysis

For cryo-EM sample grid preparation, a thin carbon film supported by a 400-mesh carbon-thickened C-flat holey grid with hole diameter 1.2 μm/spacing 1.3 μm (CF-1.2/1.3-4C, Protochips) was plasma cleaned for 2 min using Denton high-vacuum evaporator. The grid was loaded into a Vitrobot (FEI) that was preset at 100% humidity at 4°C for vitrification of our samples. 3 μl (504~900 ng in total amount) of the p53/TFIID/TFIIA/DNA assemblies was applied directly onto the grid, incubated for 2 min and then blotted for 6.5 sec prior to plunge-frozen in liquid ethane.

Cryo-EM data were collected with a JEM-2100F transmission electron microscope (JEOL) operated at 120 KeV and at a magnification of 50,000x with a defocus range of –2.5 μm to –4.5 μm. Digital micrographs were collected using a TEM 2048× 2048 CCD camera (TemCam-F224HD, TVIPS) with a pixel size of 24 μm and a calibrated magnification of 84,037× (2.86 Å/pixel) under low-dose conditions (~13 e^-^ / Å) via the semi-automatic image acquisition SerialEM software (60). Digital micrographs showed homogeneous particles in terms of lack of aggregation and apparent integrity of the intact complexes.

Particles were manually selected using boxer (EMAN) (59) for an initial total of 32,275 particles from the p53/TFIID/TFIIA/*bax* DNA dataset and 22,070 particles from the p53/TFIID/TFIIA/*hdm2* DNA dataset. The particles were then phase flipped using CTF-estimated values determined by CTFFIND3(61) and extracted using SPIDER (<u>S</u>ystem for <u>P</u>rocessing of <u>I</u>mage <u>D</u>ata from <u>E</u>lectron microscopy and <u>R</u>elated fields (62)) to a particle window size of 128 × 128 pixels (5.72 Å/pixel), following the protocol as described in (26). The particles were then normalized prior to the following data analyses. Reference-free 2D class averaging image analysis was performed using IMAGIC (63) following the procedure precisely as described (20). Using these 2D class averages as templates, we automatically picked particles from a larger cryo-EM dataset using Signature (64) embedded in the Appion image processing platform (65). A final total of 59,632 particles for the p53/TFIID/TFIIA/*bax* DNA co-complex was obtained, normalized, and low-pass band filtered to perform the 3D reconstruction using SPIDER’s multi-reference projection matching approach as previously described (29). Both rearranged and canonical forms of TFIID/TFIIA/SCP DNA from Eletron Microscopy Data Bank (accession number 10832 and 10834, respectively) were filtered to 60 Å as a reference for the first cycle of projection matching. Different initial reference models including TFIID alone, p53-bound TFIID or TFIIA-bound TFIID were also tested for model bias refinement. New 3D volumes reconstructed from the data set were used as references for all subsequent cycles of alignment (angular step sizes ranging from an initial 20° to a final 2°). The resolution throughout the refinement was determined by the 0.5 cut-off in the Fourier shell correlation (FSC) curve. The 3D structure of the p53/TFIID/TFIIA/bax assembly was filtered at a final resolution ~35 Å. All the 3D reconstructions were represented as isodensity surfaces using the UCSF Chimera package (66) developed by the Resource for Biocomputing, Visualization, and Informatics at the University of California, San Francisco (supported by NIH NIGMS P41-GM103311).

## Acknowledgement

We specifically thank R. Tjian, S. Chu, and E. Nogales for their initial support of this study, S. Zheng for generating TAF4 mAb supernatant and HeLa cells, D. King for providing the TAF4 mAb elution peptide, A. Revyakin, A. Pertsinidis and S.R. Park for their help on initial stages of the single molecule microscopy, and R. Henderson for his technical advice critical for the cryo-EM work. We also thank S.M. Shenoy, J. Hargitai, J. Wang, S. Lenny, J. Greally and E. Eng for high performance computing cluster and cryo-EM technical support. We appreciate assistance from Einstein’s AIF facility, especially F. P. Macaluso, L. Cummins and G. S. Perumal. We are grateful to M. Keogh, Charles Kenworthy and E. Nogales for critical comments of the manuscript. W. Liu is an affiliated member of the New York Structural Biology Center. M. Cianfrocco is an HHMI fellow of the Damon Runyon Cancer Research Foundation.

## References

1 MenendezD, IngaA, ResnickMA. 2009. The expanding universe of p53 targets. Nat Rev Cancer 9:724–737.

2 BiegingKT, MelloSS, AttardiLD. 2014. Unravelling mechanisms of p53-mediatedtumour suppression. Nat Rev Cancer 14:359–370.

3 TafviziA, HuangF, FershtAR, MirnyLA, van OijenAM. 2011. A single-molecule characterization of p53 search on DNA. Proc Natl Acad Sci U S A 108:563–568.

4 LuH, LevineAJ. 1995. Human TAFII31 protein is a transcriptional coactivator of the p53 protein. Proc Natl Acad Sci U S A 92:5154–5158.

5 ThutCJ, ChenJL, KlemmR, TjianR. 1995. p53 transcriptional activation mediated by coactivators TAFII40 and TAFII60. Science 267:100–104.

6 BuschmannT, LinY, AithmittiN, FuchsSY, LuH, Resnick-SilvermanL, ManfrediJJ, RonaiZ, WuX. 2001. Stabilization and activation of p53 by the coactivator protein TAFII31. J Biol Chem 276:13852–13857.

7 LiAG, PilusoLG, CaiX, GaddBJ, LadurnerAG, LiuX. 2007. An acetylation switch in p53 mediates holo-TFIID recruitment. Mol Cell 28:408–421.

8 Juven-GershonT, HsuJY, KadonagaJT. 2006. Perspectives on the RNA polymerase II core promoter. Biochem Soc Trans 34:1047–1050.

9 ColganJ, ManleyJL. 1992. TFIID can be rate limiting in vivo for TATA-containing, but not TATA-lacking, RNA polymerase II promoters. Genes Dev 6:304–315.

10 ChiT, CareyM. 1996. Assembly of the isomerized TFIIA--TFIID--TATA ternary complex is necessary and sufficient for gene activation. Genes Dev 10:2540–2550.

11 WangW, GrallaJD, CareyM 1992. The acidic activator GAL4-AH can stimulate polymerase II transcription by promoting assembly of a closed complex requiring TFIID and TFIIA. Genes Dev 6:1716–1727.

12 WhiteJ, BrouC, WuJ, LutzY, MoncollinV, ChambonP. 1992. The acidic transcriptional activator GAL-VP16 acts on preformed template-committed complexes. Embo J 11:2229–2240.

13 ChiT, CareyM. 1993. The ZEBRA activation domain: modular organization and mechanism of action. Mol Cell Biol 13:7045–7055.

14 LiebermanPM, BerkAJ. 1994. A mechanism for TAFs in transcriptional activation: activation domain enhancement of TFIID-TFIIA--promoter DNA complex formation. Genes Dev 8:995–1006.

15 ChiT, LiebermanP, EllwoodK, CareyM. 1995. A general mechanism for transcriptional synergy by eukaryotic activators. Nature 377:254–257.

16 KobayashiN, BoyerTG, BerkAJ. 1995. A class of activation domains interacts directly with TFIIA and stimulates TFIIA-TFIID-promoter complex assembly. Mol Cell Biol 15:6465–6473.

17 RheeHS, PughBF. 2012. Genome-wide structure and organization of eukaryotic pre-initiation complexes. Nature 483:295–301.

18 LiebermanPM, OzerJ, GurselDB. 1997. Requirement for transcription factor IIA (TFIIA)-TFIID recruitment by an activator depends on promoter structure and template competition. Mol Cell Biol 17:6624–6632.

19 XingJ, SheppardHM, CorneillieSI, LiuX 2001. p53 Stimulates TFIID-TFIIA-promoter complex assembly, and p53-T antigen complex inhibits TATA binding protein-TATA interaction. Mol Cell Biol 21:3652–3661.

20 LiuWL, ColemanRA, MaE, GrobP, YangJL, ZhangY, DaileyG, NogalesE, TjianR 2009. Structures of three distinct activator-TFIID complexes. Genes Dev 23:1510–1521.

21 RevyakinA, ZhangZ, ColemanRA, LiY, InouyeC, LucasJK, ParkSR, ChuS, TjianR 2012. Transcription initiation by human RNA polymerase II visualized at single-molecule resolution. Genes Dev 26:1691–1702.

22 ZhangZ, EnglishBP, GrimmJB, KazaneSA, HuW, TsaiA, InouyeC, YouC, PiehlerJ, SchultzPG, LavisLD, RevyakinA, TjianR 2016. Rapid dynamics of general transcription factor TFIIB binding during preinitiation complex assembly revealed by single-molecule analysis. Genes Dev 30:2106–2118.

23 TantaleK, MuellerF, Kozulic-PirherA, LesneA, VictorJM, RobertMC, CapoziS, ChouaibR, BackerV, Mateos-LangerakJ, DarzacqX, ZimmerC, BasyukE, BertrandE 2016. A single-molecule view of transcription reveals convoys of RNA polymerases and multi-scale bursting. Nat Commun 7:12248.

24 LiuWL, ColemanRA, GrobP, KingDS, FlorensL, WashburnMP, GelesKG, YangJL, RameyV, NogalesE, TjianR 2008. Structural changes in TAF4b-TFIID correlate with promoter selectivity. Mol Cell 29:81–91.

25 PapaiG, TripathiMK, RuhlmannC, LayerJH, WeilPA, SchultzP 2010. TFIIA and the transactivator Rap1 cooperate to commit TFIID for transcription initiation. Nature 465:956–960.

26 CianfroccoMA, KassavetisGA, GrobP, FangJ, Juven-GershonT, KadonagaJT, NogalesE 2013. Human TFIID binds to core promoter DNA in a reorganized structural state. Cell 152:120–131.

27 LouderRK, HeY, Lopez-BlancoJR, FangJ, ChaconP, NogalesE 2016. Structure of promoter-bound TFIID and model of human pre-initiation complex assembly. Nature 531:604–609.

28 AndelF, 3rd, LadurnerAG, InouyeC, TjianR, NogalesE 1999. Three-dimensional structure of the human TFIID-IIA-IIB complex. Science 286:2153–2156.

29 GrobP, CruseMJ, InouyeC, PerisM, PenczekPA, TjianR, NogalesE 2006. Cryo-electron microscopy studies of human TFIID: conformational breathing in the integration of gene regulatory cues. Structure 14:511–520.

30 BourdonJC, Deguin-ChambonV, LelongJC, DessenP, MayP, DebuireB, MayE 1997. Further characterisation of the p53 responsive element--identification of new candidate genes for trans-activation by p53. Oncogene 14:85–94.

31 Juven-GershonT, ChengS, KadonagaJT 2006. Rational design of a super core promoter that enhances gene expression. Nat Methods 3:917–922.

32 ChangGS, ChenXA, ParkB, RheeHS, LiP, HanKH, MishraT, Chan-SalisKY, LiY, HardisonRC, WangY, PughBF 2014. A Comprehensive and High-Resolution Genome-wide Response of p53 to Stress. Cell Rep 8:514–527.

33 GalasinskiSK, LivelyTN, GrebeD, BarronA, GoodrichJA 2000. Acetyl coenzyme A stimulates RNA polymerase II transcription and promoter binding by transcription factor IID in the absence of histones. Mol Cell Biol 20:1923–1930.

34 ZhangZ, BoskovicZ, HussainMM, HuW, InouyeC, KimHJ, AboleAK, DoudMK, LewisTA, KoehlerAN, SchreiberSL, TjianR 2015. Chemical perturbation of an intrinsically disordered region of TFIID distinguishes two modes of transcription initiation. Elife 4.

35 MazurSJ, SakaguchiK, AppellaE, WangXW, HarrisCC, BohrVA 1999. Preferential binding of tumor suppressor p53 to positively or negatively supercoiled DNA involves the C-terminal domain. J Mol Biol 292:241–249.

36 WeinbergRL, FreundSM, VeprintsevDB, BycroftM, FershtAR 2004. Regulation of DNA binding of p53 by its C-terminal domain. J Mol Biol 342:801–811.

37 EmamzadahS, TropiaL, VincentiI, FalquetB, HalazonetisTD 2014. Reversal of the DNA-binding-induced loop L1 conformational switch in an engineered human p53 protein. J Mol Biol 426:936–944.

38 FloydDL, HarrisonSC, van OijenAM 2010. Analysis of kinetic intermediates in single-particle dwell-time distributions. Biophys J 99:360–366.

39 MiyashitaT, ReedJC 1995. Tumor suppressor p53 is a direct transcriptional activator of the human bax gene. Cell 80:293–299.

40 PhelpsM, DarleyM, PrimroseJN, BlaydesJP 2003. p53-independent activation of the hdm2-P2 promoter through multiple transcription factor response elements results in elevated hdm2 expression in estrogen receptor alpha-positive breast cancer cells. Cancer Res 63:2616–2623.

41 LiuD, IshimaR, TongKI, BagbyS, KokuboT, MuhandiramDR, KayLE, NakataniY, IkuraM 1998. Solution structure of a TBP-TAF(II)230 complex: protein mimicry of the minor groove surface of the TATA box unwound by TBP. Cell 94:573–583.

42 AnandapadamanabanM, AndresenC, HelanderS, OhyamaY, SiponenMI, LundstromP, KokuboT, IkuraM, MocheM, SunnerhagenM 2013. High-resolution structure of TBP with TAF1 reveals anchoring patterns in transcriptional regulation. Nat Struct Mol Biol 20:1008–1014.

43 KokuboT, GongDW, YamashitaS, HorikoshiM, RoederRG, NakataniY 1993. Drosophila 230.kD TFIID subunit, a functional homolog of the human cell cycle gene product, negatively regulates DNA binding of the TATA box-binding subunit of TFIID. Genes Dev 7:1033–1046.

44 KokuboT, YamashitaS, HorikoshiM, RoederRG, NakataniY 1994. Interaction between the N-terminal domain of the 230.kDa subunit and the TATA box-binding subunit of TFIID negatively regulates TATA-box binding. Proc Natl Acad Sci U S A 91:3520–3524.

45 LivelyTN, FergusonHA, GalasinskiSK, SetoAG, GoodrichJA 2001. c-Jun binds the N terminus of human TAF(II)250 to derepress RNA polymerase II transcription in vitro. J Biol Chem 276:25582–25588.

46 LivelyTN, NguyenTN, GalasinskiSK, GoodrichJA 2004. The basic leucine zipper domain of c-Jun functions in transcriptional activation through interaction with the N terminus of human TATA-binding protein-associated factor-1 (human TAF(II)250). J Biol Chem 279:26257–26265.

47 KokuboT, SwansonMJ, NishikawaJI, HinnebuschAG, NakataniY 1998. The yeast TAF145 inhibitory domain and TFIIA competitively bind to TATA-binding protein. Mol Cell Biol 18:1003–1012.

48 OzerJ, MitsourasK, ZerbyD, CareyM, LiebermanPM 1998. Transcription factor IIA derepresses TATA-binding protein (TBP)-associated factor inhibition of TBP-DNA binding. J Biol Chem 273:14293–14300.

49 XiaoH, PearsonA, CoulombeB, TruantR, ZhangS, RegierJL, TriezenbergSJ, ReinbergD, FloresO, InglesCJet al.1994. Binding of basal transcription factor TFIIH to the acidic activation domains of VP16 and p53. Mol Cell Biol 14:7013–7024.

50 GazitK, MoshonovS, ElfakessR, SharonM, MengusG, DavidsonI, DiksteinR 2009. TAF4/4b × TAF12 displays a unique mode of DNA binding and is required for core promoter function of a subset of genes. J Biol Chem 284:26286–26296.

51 LiuX, BerkAJ 1995. Reversal of in vitro p53 squelching by both TFIIB and TFIID. Mol Cell Biol 15:6474–6478.

52 EspinosaJM, VerdunRE, EmersonBM 2003. p53 functions through stress-and promoter-specific recruitment of transcription initiation components before and after DNA damage. Mol Cell 12:1015–1027.

53 SinghSK, QiaoZ, SongL, JaniV, RiceW, EngE, ColemanRA, LiuWL 2016. Structural visualization of the p53/RNA polymerase II assembly. Genes Dev 30:2527–2537.

54 OkudaM, NishimuraY 2015. Real-time and simultaneous monitoring of the phosphorylation and enhanced interaction of p53 and XPC acidic domains with the TFIIH p62 subunit. Oncogenesis 4:e150.

55 LeveillardT, AnderaL, BissonnetteN, SchaefferL, BraccoL, EglyJM WasylykB.1996. Functional interactions between p53 and the TFIIH complex are affected by tumour-associated mutations. EMBO J 15:1615–1624.

56 PooreyK, ViswanathanR, CarverMN, KarpovaTS, CirimotichSM, McNallyJG, BekiranovS, AubleDT 2013. Measuring chromatin interaction dynamics on the second time scale at single-copy genes. Science 342:369–372.

57 HuangB, JonesSA, BrandenburgB, ZhuangX 2008. Whole-cell 3D STORM reveals interactions between cellular structures with nanometer-scale resolution. Nat Methods 5:1047–1052.

58 ChenJ, ZhangZ, LiL, ChenBC, RevyakinA, HajjB, LegantW, DahanM, LionnetT, BetzigE, TjianR, LiuZ 2014. Single-molecule dynamics of enhanceosome assembly in embryonic stem cells. Cell 156:1274–1285.

59 LudtkeSJ, BaldwinPR, ChiuW 1999. EMAN: semiautomated software for high-resolution single-particle reconstructions. J Struct Biol 128:82–97.

60 MastronardeDN 2005. Automated electron microscope tomography using robust prediction of specimen movements. J Struct Biol 152:36–51.

61 MindellJA, GrigorieffN 2003. Accurate determination of local defocus and specimen tilt in electron microscopy. J Struct Biol 142:334–347.

62 ShaikhTR, GaoH, BaxterWT, AsturiasFJ, BoissetN, LeithA, FrankJ 2008. SPIDER image processing for single-particle reconstruction of biological macromolecules from electron micrographs. Nat Protoc 3:1941–1974.

63 van HeelM, HarauzG, OrlovaEV, SchmidtR, SchatzM 1996. A new generation of the IMAGIC image processing system. J Struct Biol 116:17–24.

64 ChenJZ, GrigorieffN 2007. SIGNATURE: a single-particle selection system for molecular electron microscopy. J Struct Biol 157:168–173.

65 LanderGC, StaggSM, VossNR, ChengA, FellmannD, PulokasJ, YoshiokaC, IrvingC, MulderA, LauPW, LyumkisD, PotterCS, CarragherB 2009. Appion: an integrated, database-driven pipeline to facilitate EM image processing. J Struct Biol 166:95–102.

66 PettersenEF, GoddardTD, HuangCC, CouchGS, GreenblattDM, MengEC, FerrinTE 2004. UCSF Chimera--a visualization system for exploratory research and analysis. J Comput Chem 25:1605–1612.

